# Dwarf-shrubs dynamics in Mediterranean high mountain ecosystems

**DOI:** 10.1101/2021.10.06.463306

**Authors:** Andrea De Toma, Marta Carboni, Manuele Bazzichetto, Marco Malavasi, Maurizio Cutini

## Abstract

**Question:** Vegetation in the alpine and treeline ecotone faces changes in both climate and land use. Shrub encroachment is considered an effect of these changes, but it’s still unclear how this effect is mediated by environmental heterogeneity. Our goal is to determine which environmental factors shape the fine-scale spatial distribution and temporal trends of alpine dwarf shrub.

**Location:** Three sites in the Central Apennine, Italy.

**Methods:** We used a comprehensive set of environmental factors across a broad temporal span to model, at a fine-scale, both (1) the current spatial distribution and (2) the change in shrub cover over the past 60 years.

**Results:** Our results show that dwarf shrubs have generally increased in our study sites over the past 60 years, yet their distribution is strongly shaped by the joint influence of the fine-scale topography, productivity, land use and micro-climate. In particular, shrubs have been locally favored in areas with harsher alpine environmental constraints and stronger resource limitation. Instead, contrary to expectations, at this fine scale, warmer temperatures and the decline in grazing have not favored shrub encroachment.

**Conclusion:** Dwarf shrubs appear as a stress-tolerant, pioneer vegetation that is currently distributed mainly over areas that are otherwise sparsely vegetated. It appears that shrubs exhibit poor competitive ability to invade grasslands and, though they have increased overall, they remain restricted to the least productive areas. Fine-scale environmental heterogeneity may strongly influence future responses of dwarf shrubs in changing alpine ecosystems.

## Introduction

In the last two decades, a growing body of literature has investigated the factors driving the distribution and temporal trends of alpine shrubs (e.g. Myers-Smith et al. 2011). In particular, the existing literature focuses on shrub encroachment, which is defined as the trend of shrubs increase in density and cover at high elevation (Auken 2009). In fact, the related “greening” dynamics (i.e. the increase in productivity related to upslope migration of vascular plants, and in particular of woody species; Carlson et al. 2017) has been observed worldwide in alpine and arctic ecosystems and is often ascribed to climate warming (e.g. Hudson et al. 2011; Elmendorf et al. 2012; Ylänne et al. 2015; Francon et al. 2017; Myers-Smith & Hik 2018; García Criado et al. 2020). A significant increase in shrub cover in these ecosystems has the potential to dramatically alter the microclimate, nutrient cycling, and species composition (Myers-Smith et al. 2011). Despite the growing attention to shrub encroachment (e.g. Dullinger et al. 2003; Tape et al. 2006; Komac et al. 2013; Francon et al. 2017), the factors shaping the local distribution and temporal trends of shrubs in arctic and alpine areas remain poorly understood (Hallinger et al. 2010). Indeed, divergent temporal trends have been so far observed for dwarf and tall shrubs (Elmendorf et al. 2012): while tall, woody and deciduous shrub species generally follow the much-described encroachment trend, the dynamics of dwarf shrubs is less consistent, often showing evidence of exposure to increased physiological stress on mountain summits or in Arctic regions. In some areas, this can even result in a significant decline of their representation (García et al. 1999; García 2007; Wipf et al. 2009; Wheeler et al. 2014; Pellizzari 2014; Pellizzari et al. 2017; Gamm et al. 2020; Buchwal et al. 2020). Dwarf shrubs are evergreen growth-forms occurring up to and above the treeline that grow prostrate to the ground (0.1 - 0.4 m). They typically form “cushions” to withstand the extreme environmental conditions encountered at high altitudes (Körner 2003). In this way, dwarf shrubs are potentially decoupled from the environmental conditions that drive the encroachment of taller shrubs; their growth is likely more associated with and controlled by the exposure to microclimatic and local soil and snow cover conditions (Körner 2012).

Thus, in order to understand the ongoing dwarf shrub dynamics, it is important to consider multiple potential drivers simultaneously and at a fine scale. Yet, such a comprehensive approach is still rare (Filippa et al. 2019). Indeed, most studies investigating the distribution and temporal trends of alpine dwarf shrubs took into account only a few factors, mainly focusing on the individual influence of either climate change or land-use abandonment (mainly, the abandonment of traditional grazing practices; Bjorkman et al. 2020). Moreover, these studies largely overlooked the effect of other fine-scale drivers of shrub distribution such as local soil conditions related to topography and productivity, snow cover and microclimate, and, above all, the interactions existing among these factors in alpine environmental mosaic (Myers-Smith 2020). Disregarding these has so far prevented comprehensive understanding of the true drivers of dwarf-shrub dynamics and the potential for encroachment in alpine and arctic regions. Also, as dwarf shrubs are long-lived and clonal species, the analysis of their encroachment dynamics should focus on a wide temporal span (i.e. from decades to centuries), which further complicates their investigation (Myers-Smith et al. 2015).

Among dwarf-shrub species, *Juniperus communis* var. *saxatilis* (also known as *J. communis* subsp. *nana* or subsp. *alpine, sensu* World Check-List of Selected Plant Families, hereafter referred to as *J. communis*) occurs throughout the northern hemisphere (Farjon 2010). It is a light demanding, drought and frost tolerant species adapted to the alpine and arctic climate. In the past decades, the temporal trends of *J. communis* have greatly varied across the northern hemisphere: on British islands and in Spanish mountains, populations have declined (Ward 1977; García et al. 1999; Verheyen et al. 2009; Broome & Holl 2017), whereas in Greenland and southern Mediterranean, *J. communis* has shown an encroachment trend (Rosen & Barthlott 1991; Pellizzari et al. 2017). This broad variation, which appears largely unrelated to the latitudinal (i.e. climatic) gradient, suggests that local factors are likely to be important in determining the fine-scale spatial distribution of this dwarf-shrub species, ultimately influencing its overall regional increase or decrease (Tumajer et al., 2021). Thus, *J. communis* represents an ideal species for studying how dwarf-shrubs respond to fine-scale variation in soil condition, micro-climate, and land use (Carrer et al. 2019).

To improve our understanding of dwarf shrubs-related processes, it is thus crucial to: (i) take into account the simultaneous influence of multiple fine-scale distribution drivers; (ii) use a set of environmental variables derived at a fine resolution to perform analyses at the local spatial scale; and (iii) consider a wide temporal scale (e.g. using data on present and past dwarf shrubs distribution).

In view of this, here we aim at analyzing fine-scale temporal trends and current local patterns of *J. communis* in three Mediterranean alpine areas in the Central Apennines. Specifically, we: (i) investigate whether the distribution of *J. communis* has generally increased or decreased over the last 60 years; (ii) explore what shapes the current local distribution of *J. communis*; (iii) identify from a comprehensive set of land-use, climate, topographical and productivity factors those that have locally driven the potential encroachment or decline observed over the last 60-yrs;

The reason we focus on the Mediterranean mountains is that they preserve fragments of alpine summit vegetation within a peculiar semi-arid climate and land-use context. These climatic conditions have progressively exacerbated over the past decades as the Mediterranean area is particularly affected by climate change, which is 20% faster in this region than in the rest of the globe (Giorgi & Lionello 2008).

## 2. Material and methods

### 2.1. Study sites

We conducted our study in the alpine zone and treeline-ecotone of three limestone massifs located in the Central Apennines: Mt. Terminillo, Mt. Duchessa, and Mt. Ernici (Fig 1). These massifs occur along a latitudinal gradient that allows capturing the variation in both land-use trends and climate features in our study system. The Apennines stretch across the Italian peninsula and, due to their proximity to the Mediterranean Sea, represent a transition from continental to sub-Mediterranean climate features, with warm and dry summer seasons (Cutini et al. 2021). The mean annual temperature in our study sites ranges from 6.4° to 4.5° and the mean annual rainfall from 1790 mm to 738 mm (for Mt. Ernici and Mt. Terminillo respectively). In summers, lack of water may occur here as a combined result of the climate, topography, calcareous lithotype, and historical land-use that has led to thin and immature soil. Regarding land-use, though the Central Apennines have historically been characterized by grazing, this activity has undergone a major decline after the World War II due to the socio-economic transformations (appendix S1). Despite this general trend, the three study sites show at present areas where grazing has been abandoned as well as those where it still persists. Human activities, together with natural environmental heterogeneity, lead to a finely grained vegetation matrix intermingled with scattered woody-dwarf-shrublands, which are nearly exclusively dominated by *J. communis* in a prostrate form.

**Figure 1.**
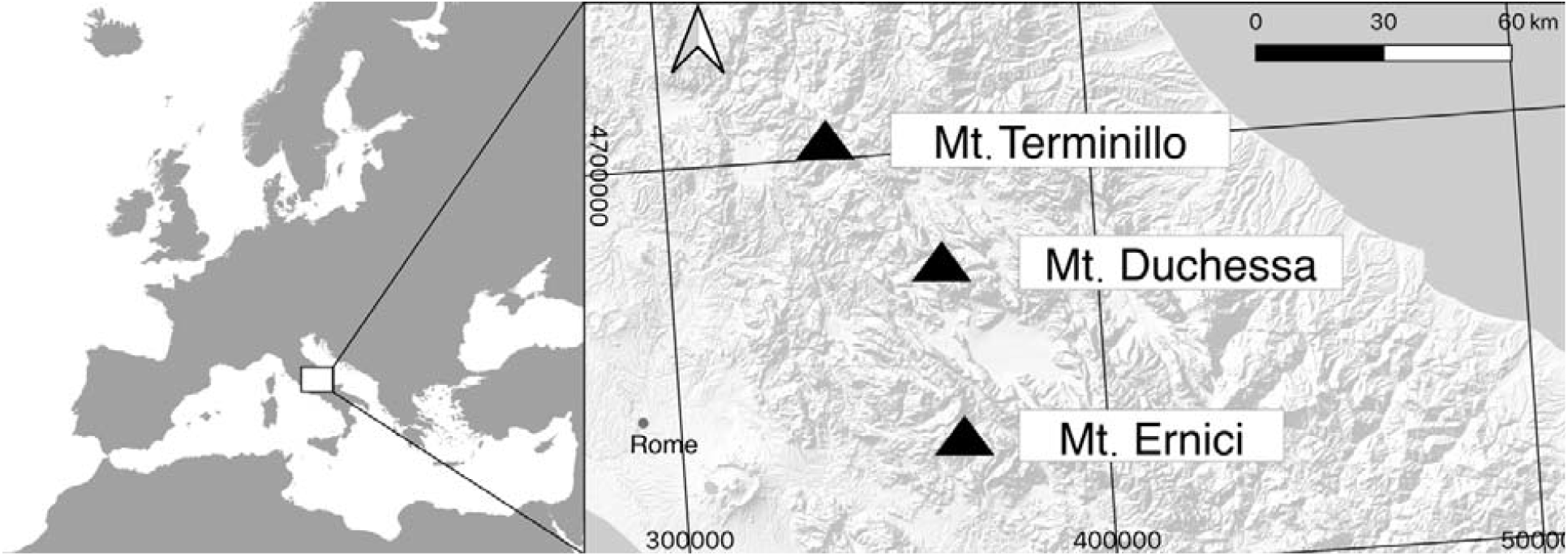
Map of Europe (scale 1:30,000,000, CSR: WGS-84 / UTM zone: 33N) with focus on Central Italy (1:300,000) showing the three study sites. Mt. Terminillo (E335689, N470480), Mt. Duchessa (E362889, N4671881) and Mt. Ernici (E364166, N4631951).

### 2.2 Sampling design

First, in a Qgis 3.6 environment (QGIS.org, 2019), we defined the boundaries of each study site by mapping the current treelines (1:900 resolution) using Google Satellite photos from 2012. Following Körner (2003), we identified the alpine zone as the areas occurring above the treeline. For brevity, along the text we will use alpine to include alpine and subalpine space, i.e. also including the treeline ecotone. We defined the treeline by connecting the highest patches of forests through photo interpretation. This led us to focus on a total area of 33,557 km^2^, with a mean elevation of 1845 m a.s.l. Second, we used a 5m resolution digital terrain model (DTM) derived from a regional technical map to stratify our random sampling points by slope. This was done due to the relevance of the slope to many factors (such as moisture, snow, soil accumulation) and to avoid the underrepresentation of flat areas. We obtained three different classes of slopes (0-15%; 15-30%; 30-70%). Then, we randomly generated up to 100 points for each class in each site (or, where less than 100 points were available, the maximum possible number of points), thereby obtaining 860 sampling points. Around each point, we created a circular buffer with a diameter of 100 m. Finally, within each 100 m-buffer, we mapped the present (2012) and past (1954) dwarf-shrub patches (see 2.2.1) and summarized the information on climate, snow cover, grazing, topography, and productivity (see 2.2.2).

#### 2.2.1. Present and past shrub distribution

To gather information on the present and past shrubs distributions, we mapped the dwarf-shrub patches (1:900 resolution) through photo-interpretation using Google Satellite photos from 2012 and historical aerial high resolution (1:30,000, 2400 dpi) photos from 1954 available from the IGM (Italian Geographic Military Institute). The photo interpretation of dwarf-shrubs was relatively easy thanks to their cushion form; these patches are mainly composed by *J. communis* in a prostrate form which is an evergreen, procumbent and well recognizable shrub often < 10 cm high (Thomas et al. 2007). However, tallgrass may form similar cushion structures (e.g. *Brachypodium genuense*) which can hamper accurate differentiation from perennial dwarf shrubs. To facilitate the classification, we used autumn and winter satellite photos where necessary. We mapped the coalescent dwarf-shrub patches into a single patch. Based on the recent map (2012), the other time point (1954) was processed in reverse order by editing, with information being adapted to older time levels.

Overall, we mapped over 12,500 dwarf-shrub patches (1954 and 2012). For each buffer, we derived the current and past shrub occurrence (presence/absence within the buffer); current and past shrub cover (area of patches mapped within the buffer); and the temporal change of the shrub (computed as the cover area change between the present and the past). The current shrub occurrence and cover were used to investigate the main drivers of the current fine-scale distribution (Tab. 1), while the calculated temporal change was used to 1) evaluate whether dwarf shrubs in the study area are generally encroaching or declining, and 2) identify the fine-scale drivers of these temporal trends. Moreover, to take into account the recruitment of shrubs (i.e. the process by which new individuals found a population or are added to an existing population), we calculated the encroachment of shrubs only in buffers that contained no shrub patches in 1954.

**Table 1.**
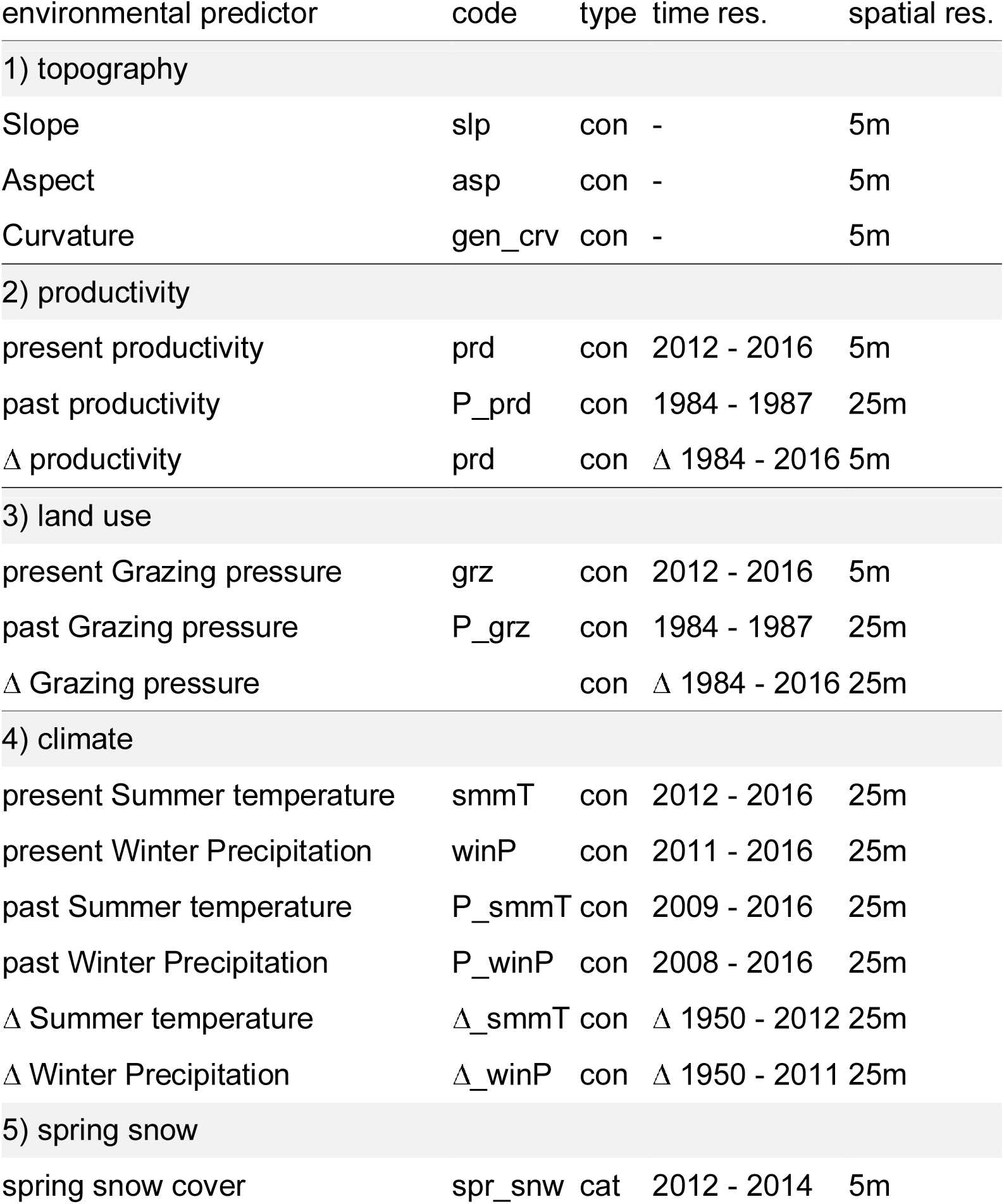
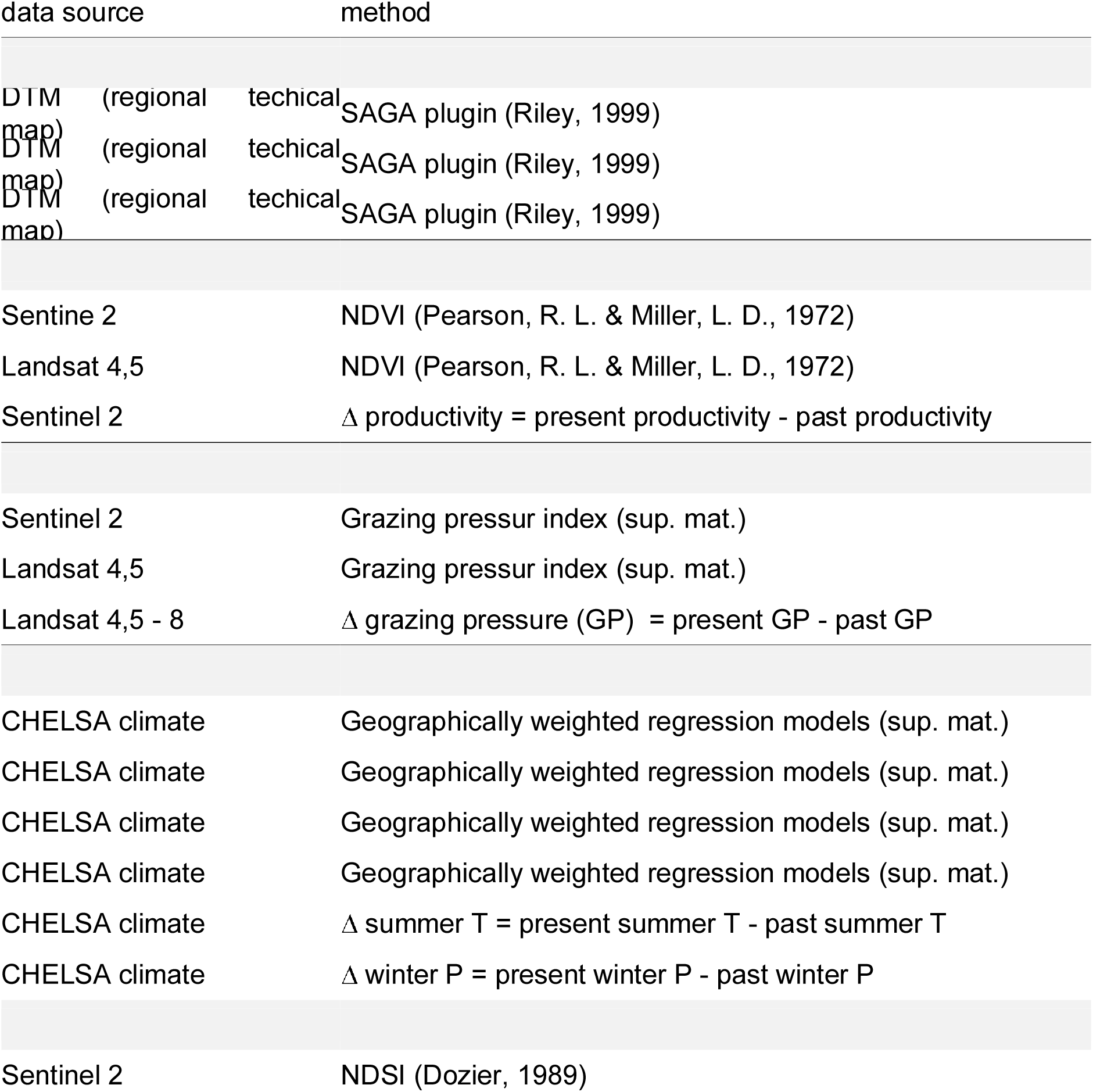
Environmental predictors used in the study, indicating the type of variable (con= continuous vs. cat= categorical), the time resolution (the time interval to which the vrible refers), the spatial resolution of the environmental layer, the data source and the method used to derive the variable (with reference where appropriate). Δ indicates a range or change in thetime interval. “Code” refers to the code used in Fig. 3 and in supplementary materials for the variable.. “Data source” refers to the primary source from which the data is derived (see details in the main text and Appendix S3)

#### 2.2.2. Drivers of shrub distribution and encroachment

To model shrub occurrence, cover and change, we derived environmental variables representative of the main drivers thought to influence shrub encroachment within each buffer and (where relevant) at the the two time points, namely grazing, climate, snow cover, topography, and productivity (Bjorkman et al. 2020).

First, we computed a **grazing pressure** index using the Normalized Difference Vegetation Index (NDVI) values calculated from multispectral images from 2012 (for the current grazing pressure) and 1984-1987 (for the past grazing pressure). Specifically, we computed grazing pressure (both in the present and in the past) as:

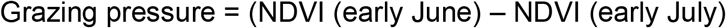

which is, therefore, defined as the difference between NDVI at the time before grazing starts and the time of an observable grazing impact on the biomass (while avoiding impact of summer drought on the biomass). We were able to use this index because of the traditional transhumance practice in our areas: shepherds usually start grazing in early June due to regional laws. For the past interval (1984-1987) we used images from Landsat 4–5 (the oldest available multispectral images for our study area), while for the present (2012) we used two sets of images: Landsat 8 (for the comparison between the present and the past) and Sentinel 2 (for the present-time models, to achieve a better resolution). We thus calculated the average grazing pressure and the difference in grazing within each buffer. To check that this grazing index derived from remotely sensed data captured grazing pressure in our study sites correctly, we also compared it with field-collected grazing proxies (Appendix S2).

Second, we gathered present (2008-2012 mean) and past (1950-1954 mean) climate data. In particular, we focused on four bioclimatic variables: (1) the average annual temperature, since low temperatures are the most characteristic factor of alpine environments; (2) the temperature of the warmest quarter since it can indicate the probability of the presence of a summer water restriction, characteristic constraint of Mediterranean areas; (3) the precipitation in the coldest quarter, part of which is snow that protects the soil and the low vegetation from the constraint of frost events; (4) the annual temperature range, in order to highlight whether the two types of environmental contraints differ in the cold and warm seasons. Because global climate layers are not available at a resolution sufficient for capturing local micro-climatic variability of mountain environments, we derived fine-scale topoclimatic layers through downscaling techniques. Specifically, we first collected climate layers at a ∼1 km resolution from the CHELSA database. Then, to account for the micro-climate heterogeneity, we carried out a 25 m-resolution downscaling using geographically weighted regression models (GWR; Appendix S3). Finally, we obtained the present and past average values of each downscaled variable within each buffer, as well as the temporal difference in climate data between 1954 and 2012.

Third, to gather information on snow cover in springtime, we computed a normalized difference snow index (NDSI) from Sentinel-2 multispectral images according to Dozier (1989). The NDSI naturally ranges from 0 to 1 with high values indicating the presence of snow. Following Dozier (1989), we transformed continuous spring NDSI into a binary variable: we classified values above 0.4 as 1 (“snow present”), and those below 0.4 as 0 (“snow absent”), respectively. We used snow cover both in late winter and spring, where early snowmelt left some areas without snow. Dwarf shrubs are known to be extremely dependent on snowmelt time (Wipf & Rixen 2016). Unfortunately, due to the frequent presence of the cloud cover in winter, satellite images suitable for calculation of past NDSI were not available. Therefore, we were unable to calculate past snowmelt patterns and its changes over time. However, we assume that the pattern of snow accumulation and melting computed from the present data can be used as a proxy for the past data as this variable is mainly affected by topography and microclimate (Tappeiner et al. 2001).

Fourth, to account for the alpine topographic heterogeneity, we obtained topographical data from a digital terrain model (DTM) at a 5 m resolution. We subsequently computed average elevation, slope, aspect, ruggedness (amount of elevation difference between adjacent cells of a DTM) and general curvature (that represents the shape or curvature of the relief, where positive values indicate upward concavity) within each buffer using SAGA plugins (Riley et al. 1999). We chose this set of variables since they describe well the accumulation, formation and erosion of soil, and more generally are good proxies for environmental micro-heterogeneity in these environments (Geitner et al. 2020).

Finally, plant productivity (hereafter: productivity) has strong implications for ecosystem services in alpine landscapes, such as erosion control through root systems (Löbmann et al. 2020) and pasture resources for livestock. To obtain information about productivity, we computed the NDVI at the peak of the growing season (i.e. late June). The NDVI is generally used as a significant predictor of soil nutrients and moisture (Löfgren et al. 2018). As for the grazing index described above, we used multispectral images from both the present (2012, Sentinel 2 and Landsat 8) and the past (Landsat 4-5). We computed the average NDVI within each buffer (computed only outside the shrub patches, to avoid circularity).

### 2.3. Models of shrub distribution and dynamics

We modelled the current occurrence and cover (distribution) of the shrubs as well as shrub encroachment separately as a function of the predictors related to grazing, climate, snow cover, topography, and productivity. Predictors were standardized (i.e., rescaled so that the mean is 0 and standard deviation corresponds to the normal distribution standards) to make them vary on a comparable scale. To reduce multicollinearity, we tested pairwise Pearson correlations among predictors, eventually retaining the least correlated (r < 0.35) and most ecologically meaningful ones. Specifically, out of the original fourteen variables, we retained seven predictors for fitting the models: winter precipitation, summer temperature, spring snow cover, grazing pressure, slope, general curvature and productivity.

To model current shrub distribution, we used a two-step approach. First, we modelled the presence/absence of shrubs within 100 m buffers using a binomial generalized linear mixed model (GLMM, function *glmer* in R, package lme4) with logit link (hereafter referred to as the “occurrence model”). Second, we fitted a linear mixed model (LMM, function *lme* in R, package lme4) assuming normally distributed errors to model the current (logit-transformed) shrub cover (hereafter referred to as “cover model”). The cover was modelled only considering buffers where shrubs were present (i.e. 678 out of the 860 buffers). Third, we modelled the temporal trends in shrub cover from 1954 to 2012, expressed as a binary variable, using a binomial GLMM (hereafter referred to as “encroachment model”). In particular, we calculated the change in cover (encroachment) as the difference in shrub cover between 2012 and 1954, divided by the cover in 1954 in order to account for how shrubs of different sizes in 1954 changed in relative terms. Then, as preliminary analyses showed that declines in cover were extremely rare (see results and Fig. 2), we transformed the ratio within each buffer into a binary variable: values of cover change above 1, indicating encroachment, were classified as 1, while values below 1, indicating absence of encroachment, were classified as 0.

**Figure 2.**
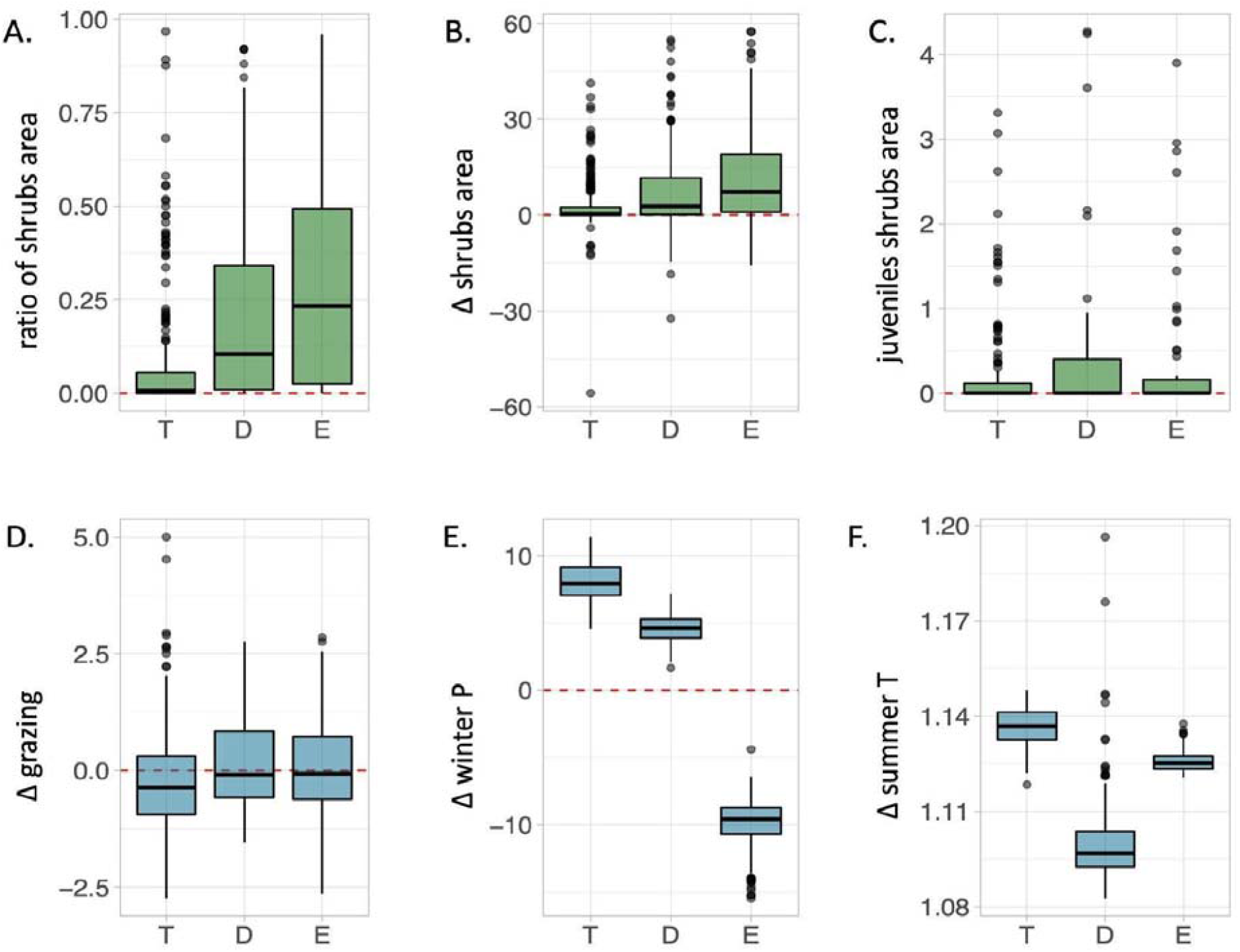
Overall distribution of shrub (green) and environmental (blue) time trends across the three sampling sites. We refer to sites as T = Mt. Terminillo, D = Mt. Duchessa, E= Mt. Ernici **A**) ratio of average shrub cover within buffers at present (2012). **B**) average shrubs cover change between the present (2012) and the past (1954). **C**) average shrub cover recruitment, taking into account the average shrubs area in the present (2012) only in the sampling buffers without shrub patches in the past (1954). **D**) average grazing pressure change between the present (2012) and the past (1986). **E**) average winter precipitation changes between the present (2012) and the past (1954). **F**) average summer T change between the present (2012) and the past (1954).

In all three models, we included all seven uncorrelated predictors as main fixed effects as well as interactions between the productivity and all other predictors. Indeed, the effect of other environmental factors on dwarf shrubs depend on productivity, e.g. high temperatures during the growing season will likely not have the same effect in highly productive sites and in poor arid soils. At the same time, productivity affects other factors, such as grazing – on a landscape scale, livestock can choose the most productive areas to graze (Adler et al. 2001). Modeling productivity interactions with other environmental factors enabled us, therefore, to take into account the moderating influence of productivity on the relationship between shrubs and other environmental features. Further, we included the site as a random intercept to account for the lack of independence among observations collected in the same area. Moreover, we included random slopes for grazing pressure and summer temperature across sites, to account for the fact that the three sites differ markedly in climate and average grazing pressure.

The encroachment model, compared to the previous two, takes into account the same set of main effect terms, interactions and random slopes, but in this case, we used the variables calculated from the historical time points. In addition, in order to take into account the degree of land-use changes and climate change relative to the past conditions, we also included the difference between the present and the past as additional terms in the case of the productivity, climatic, and grazing variables. For example, Δ summer temperature was calculated as T_present -_ T_past_, so that positive values indicate an increase in temperature over time and vice versa.

For all models, we performed backward model selection, on interactions first, using maximum likelihood (function lrtest in R, package lmtest) to compare hierarchically nested models. This test determines whether a reduced, simpler model (i.e., one formulated by constraining a given number of parameters to 0) fits the data equally well as a more complex model (where parameters are not assumed to have value 0). We evaluated the goodness-of-fit of the models computing marginal and conditional adjusted-R-square (function r.squaredGLMM in R, package MuMIn). Model assumptions were checked visually inspecting residuals plots.

We performed all statistical analyses in R, version 3.6.2 (R Core Team, 2019).

## 3. Results

### 3.1 Overall shrub distribution and temporal trends

Results showed that shrubs are currently widely distributed in our study area. In our analyses, we found shrub patches in 78.5% of our 860 sampling buffers, with heterogeneous distribution between the three sites (on average, shrub cover was greatest in the southernmost Ernici site, Fig. 2a). Regarding shrub cover change, we observed an overall increase in cover (encroachment trend) between 1954 and 2012. Wilcoxon Signed-Ranks test indicated that this increase is significantly different from zero (V=211950, p < 0.0001). We observed an increase in shrub cover in 492 of 860 plots with a average increase of 6.94% in the cover across the three sites (Fig. 2b). This cover change was correlated (Pearson’s R = 0.497) with an increase in the mean size of shrub patches within the buffers (Fig. 2c). To account not only for the expansion of already existing shrubs but also for the establishment of juvenile shrubs, we also tracked the subset of sampling buffers where patches of *J. communis* were absent in 1954. We observed *J. communis* recruitment in 11.74% of the overall 860 sampling buffers. Recruitment occurred in 30.77% of the northernmost site (Mt. Terminillo), compared to 33.87% of the southernmost site (Mt. Ernici) and with the highest rate in Mt. Duchessa (49.23%).

### 3.2. Models of shrubs distribution

The best model explained 54.45% of the total variance in shrub occurrence (18.51% marginal R^2^). Variables retained in the final model were: summer temperature, winter precipitation, spring snow cover, slope, general curvature and productivity (Fig. 3). Shrub occurrence was positively related to general curvature (Fig. 3a), indicating that shrubs tend to occur in hilly surfaces associated with runoff and thinner soils. In contrast, shrub presence is less likely in buffers where spring snow, summer temperature, winter precipitation and productivity are high. Finally, the productivity significantly interacts with summer temperature and slope in the sense of areas with high productivity being much more susceptible to the negative effect of summer temperature and the positive effect of slope.

**Figure 3.**
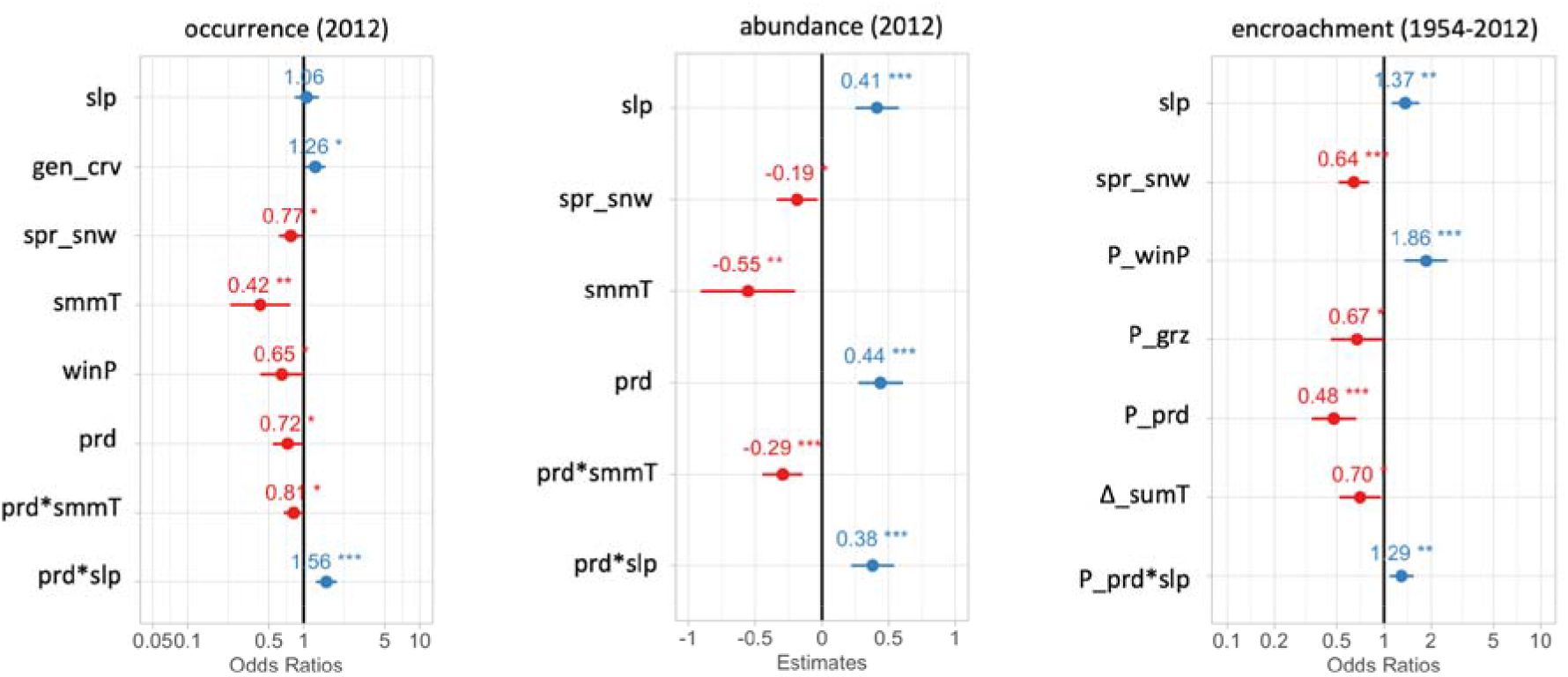
Forest plots of occurrence (a), cover (b) and encroachment (c) regression models estimates. Numbers refer to the effect size, the line to the confidence interval. In detail: **slp** = slope; **gen_crv** = general curvature; **spr_snw** = spring snow; **winP** = winter precipitation; **prd** = productivity; **P_winP** = past winter precipitation; **P_grz** = past grazing; **P_prd** = past productivity; Δ**_smmT** = difference in summer temperature.

For shrub cover, the best model explained 47.99% of the total variance (9.12% marginal R^2^), and included the same predictors as the occurrence model, except for winter precipitation and general curvature. Where shrubs were present, the cover was strongly positively related to slope and NDVI (Fig. 4b). In contrast, shrub cover decreased with the increase of summer temperature and spring snow. Finally, there was a significant interaction of NDVI with summer precipitation and slope, indicating, similar to the shrub occurrence, that the effect of temperature and slope on shrub cover was dependent on local productivity.

**Figure 4.**
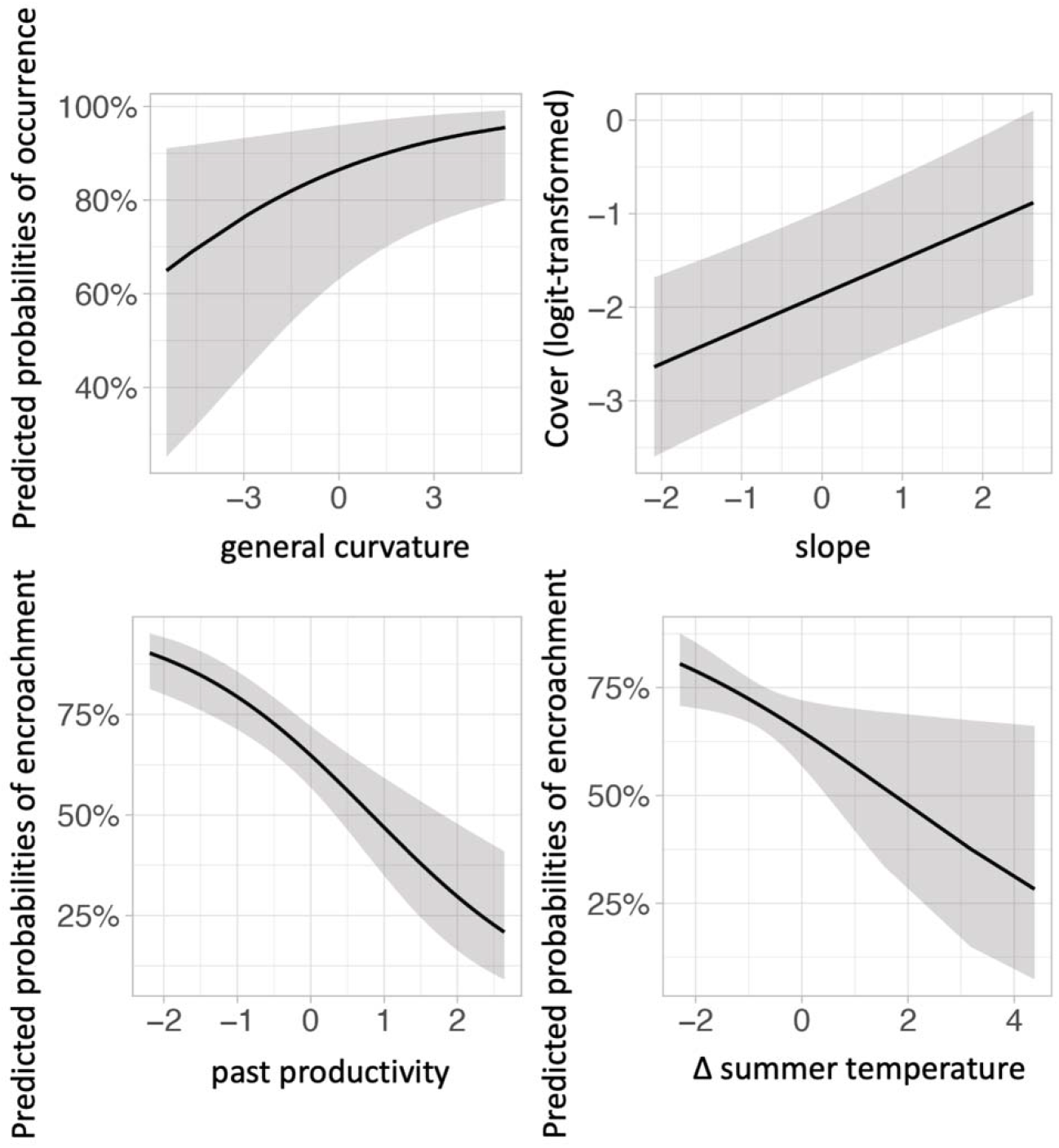
Partial effects of regression models (values of predictors on the x-axis are centered and scaled): A) relationship of the probability of occurrence with the general curvature (positive values indicate concave areas); B) relationship of probabilities of occurrence with slope (positive values indicate steeper slopes); C) relationship of probability of encroachment with past productivity (positive values indicate more productive areas); D) relationship of probability of encroachment with variation in summer temperature (positive values indicate a temperature increase (C°)).

### 3.3. Models of shrub encroachment

The final shrub encroachment model explains 25.70% of the total variance (23.98% marginal R^2^). Shrub encroachment since the 1950s in our study area is best explained by slope and snow cover, but also by past winter precipitation, grazing pressure, productivity and by the increase in summer temperature (Fig. 3). Shrub encroachment is particularly strongly positively associated with slope and past winter precipitation. By contrast, encroachment is negatively associated with spring snow cover, past grazing pressure, past productivity and increase in summer temperature (Fig. 3 c-d). Finally, there is a significant interaction of past productivity with slope, indicating that positive effects of slope are more pronounced where local productivity is higher.

## 4. Discussion

### 4.1 Overall shrub distribution and temporal trends

In our study area, dwarf-shrubs of *J. communis* represent a widely and heterogeneously distributed vegetation type. This current widespread distribution of dwarf shrubs in the alpine zone appears to be partially the result of large cover increases of *J. communis* since the 1950s in the three sites. Indeed, between 1954 and 2012, we observed an increase in shrub cover in more than half of our sampling buffers, suggesting a generalized encroachment trend of dwarf shrubs rather than decline as observed in other areas (Ward 1977; García et al. 1999; Verheyen et al. 2009; Broome & Holl 2017).

Dwarf-shrubs are more widespread in the southernmost site (Mt. Ernici) than in the other two, which is also the warmest, least productive and most grazed one. Temporal increases in the cover across sites mirrored the present distribution patterns, with the greatest increases occurring in the southernmost site (Mt. Ernici) and the lowest increases in the northernmost site (Mt. Terminillo). These findings are partially in contrast with data reported across latitudinal gradients in other Mediterranean mountains region (García et al. 1999), where higher temperatures, overgrazing and summer drought were related to declining populations of *J. communis* dwarf shrubs in the southernmost sites (Verheyen et al. 2009). García (2007) has ascribed the *J. communis* decline to a strong recruitment limitation under the summer drought stress, leading to seed abortion and mortality, and reduced germination. Contrary to García (2007), recruitment does not seem to have been limited by these factors in our sites: we observed the same high rate of recruitment (∼30-34%) in the northernmost and less grazed site and the southernmost and most grazed site. These findings add to a large set of contrasting temporal trends of dwarf shrubs in the alpine and arctic areas and suggest that, under certain conditions, encroachment by *J. communis* may occur even at the boundary of their ecological tolerances.

### 4.2 Drivers of shrubs distribution and temporal trends

The drivers behind the current fine-scale spatial distribution in *J. communis* cover were (with some exceptions) well aligned with the drivers of its temporal changes. Our models confirmed that climatic conditions are important drivers of both shrub distribution and encroachment, lending support to previous evidence of the role of the climate change as an underlying cause of shrub dynamics in alpine areas (García Criado et al. 2020). However, we showed that despite widespread encroachment of dwarf shrubs potentially linked to large-scale climate and land-use changes, fine-scale distributional changes of *J. communis* were highly dependent on topography and productivity. In fact, topographic heterogeneity as well as variation in fine-scale productivity strongly shaped shrub patterns in these Mediterranean summits, with an important modulating role of productivity on the climate effects. Surprisingly, our results suggest that spatial variation in current grazing pressure had no influence on the current dwarf-shrub distribution and encroachment.

#### 4.2.1. Grazing

Although grazing and its abandonment is considered the main driver of shrub distribution and encroachment in alpine areas (Theurillat & Guisan 2001; Dullinger et al., 2003; Kornac et al., 2013), we found its effect significant only in the shrub encroachment model out of the three constructed models (moreover, this was only true where the past grazing intensity is concerned). Surprisingly, the signals of dwarf shrub encroachment were not more significant in areas abandoned by grazing in the last decades, suggesting that although land abandonment is an important driver of encroachment at a coarser scale, it does not necessarily explain where encroachment will most likely occur at a fine scale. In fact, moderate grazing appears even to promote *J. communis* invasion a) by creating open spaces that can promote advancement and reducing competition with grasses and b) by acting as a vector for juniper propagules (Broome & Holl 2017; Broome et al. 2017)

In particular, encroachment by *J. communis* was favored in the areas that were already historically poorly grazed, while the probability of encroachment was lower in the areas that were heavily grazed in the 1980s. This might be a long-lasting consequence of grazing animals feeding on *Juniper* seedlings (Stankeva-Terziyska et al. 2020), which caused high past seedling mortality and low recruitment in these areas of which we still see the signals today. Another possible explanation is that it is a consequence of artificial shrubs removal by shepherds in the historically most favorable grazing sites.

#### 4.2.2 Climate

Contrary to our expectations, locally warmer summer conditions do not favor *J*.*communis* in the central Apennines and dwarf-shrubs did not thrive where temperatures increased the most as could be expected since climate warming has been previously pinpointed as an important driver of shrub encroachment (Myers-Smith et al. 2011). Rather, their distribution increased where warming effects were more moderate. The same lack of a relationship of encroachment with higher increase in temperatures for *Juniperus communis* was found in southwest Greenland (Trkal & Lehejček 2017, Lehejček et al., 2017). However, in our case, these alpine shrubs in Mediterranean mountains are already at the lower boundary of their geographical range and likely at the warm limits of their temperature niche (Carrer et al. 2019). Our results suggest that further exacerbation of heatwaves and drought events, as expected in climate change scenarios especially in Mediterranean mountains, could eventually result in future generalized mortality and loss of these shrubs, inverting the past encroachment trends. Although adult individuals are highly drought-resistant, the reproduction of *J. communis* seems to be particularly vulnerable to drought events and heat waves (García et al. 1999; Verheyen et al. 2009; Gruwez et al. 2013; Gruwez et al.. For this reason, even though moderate temperature increases seem to have led to the widespread *J. communis* encroachment in our sites, conditions where temperatures have become too extreme were apparently not suitable for new colonization and recruitment.

Another somewhat surprising result related to climate was that *J. communis* distribution was negatively related to spring snow cover and winter precipitation. In alpine and arctic biomes, winter snow cover and late (spring) snowmelt can be beneficial since this combination results in high soil moisture that can be capitalized by plants during the next growing season (Winkler et al. 2018) while offering protection from winter desiccation and freezing events that can affect plants (Sturm et al. 2001; Sturm 2005; Hallinger et al. 2010; Carlson et al. 2015; Francon et al. 2020) as well as the soil microbial community (Körner 2003). However, this advantage is not consistent across shrub species (Wheeler et al. 2014). In our study, *J. communis* distribution was indeed negatively related to spring snow cover and winter precipitation, as also reported previously by Carrer (2019). Although these may represent a benefit at the first sight, snow cover also prevents photosynthetic activity of dwarf shrubs; thus, late spring snowmelt results in a delay in the beginning of the growing season (Hallinger et al. 2010; Francon et al. 2017; Pellizzari et al. 2017) and the advantage offered by snow cover seems to be counterbalanced by a shorter growing season (García-Cervigón et al., 2018). In addition, *J. communis’* extreme tolerance to winter drought stress (Mayr et al. 2010) allows it to inhabit snow-free areas and capitalize on the benefits of a longer growing season. Suillivan’s 2001 study (cited in Thomas et al. 2007) found a high mortality rate of *J. communis* under prolonged snow cover.

More in detail, in contrast to what we found for the current occurrence patterns of *J. communis*, past winter precipitation seems to be a factor that favored the probability of local shrub encroachment over time. A possible explanation is that *J*.*communis* juveniles, the most fragile stages of the life cycle (Broome & Holl 2017), may have required snow protection at least in the winter. This could explain why today we observe cover increases where these juveniles best survived in the past. Also, areas with both higher winter precipitation and early snowmelt may be affected by frequent freeze/thaw cycles (Freppaz et al. 2007) which tend to increase the amount of available nutrients. This might represent an additional mechanism through which early colonization by shrubs was favored in these conditions. However, snow-cover in spring had also a negative effect on encroachment, confirming that the positive effects of a longer growing season are overall greater than those of snow protection for this hardy species.

#### 4.2.3. Topography and productivity

Besides climatic factors, topography was another important driver of shrub distribution and dynamics. In particular, the occurrence, cover and encroachment of shrubs were greater in steeper conditions and in summit areas (positive, i.e. concave, curvature). This finding is in line with previous evidence suggesting a strong ability of *J. communis* to colonize harsh environments and infertile soils (Ward 1977). Indeed, higher slopes and positive curvatures lead to increased erosion and wind desiccation (summit effect; Körner 2012) that negatively affect soil formation, accumulation and persistence of snow as well as the accumulation of surface runoff water (Geitner et al. 2021). Nonetheless, *J. communis* is a typical shrub species of poor soils and harsh environments (Thomas et al. 2007) and, due to its cushion form, can intercept runoff water, reducing soil erosion and wind desiccation under its canopy. In this way, *J. communis* can create “islands of fertility” under its canopy (DeLuca & Zackrisson 2007; Allegrezza et al. 2016) and decouple itself from local soil conditions, thereby, once established, colonizing unfertile areas with higher slope and curvature.

At the same time, locally high productivity generally reduced shrub occurrence and encroachment, especially in warmer areas. The conservative strategy of *J. communis* and its poor competitive ability (it suffers from the lack of light; Niinemets & Valladares 2010) probably drives it to colonize areas where competition with other species is lower. Indeed, the negative relation of shrub occurrence with productivity may support the hypothesis that *J. communis*, like other light-demanding dwarf shrubs (e.g. *Pinus mugo*), may suffer if shaded by dense grassland, especially in the early life stages (Dullinger et al. 2003). Coherently, according to Broome & Holl (2017) and Broome et al. (2017), *J. communis* may benefit from open space. Additionally, other studies support the hypothesis that these aboveground patterns reflect belowground competition between herbaceous species and shrubs based on their different root architecture systems. Indeed, Rosén (1988) found strong belowground competition for water resources between *J. communis* individuals as well as between *J. communis* and other woody and non-woody species. In particular, shrubs may suffer from the better ability of shallow root systems of herbaceous plants to capitalize on the spring and summer soil moisture (Morris et al. 2016) or the mechanical resistance to the invasion of grassland root systems (Abbate et al. 1994). Moreover, *J. communis* may remain relatively unaffected by poor soil productivity thanks to its ability to obtain nitrogen and phosphorus from symbiosis with feather mosses (Houle & Babeux 1994; DeLuca & Zackrisson 2007), especially on the lateral branches. However, where shrubs are able to survive competition with the surrounding herbaceous vegetation at the juvenile stage (i.e. where they can establish and grow to maturity), soil productivity actually favors juniper growth to cover greater areas (positive effect of productivity on shrub cover in the cover model).

## 5. Conclusion

The overall goals of this work were to investigate the spatial and temporal patterns of a widespread and characteristic alpine dwarf shrub in Mediterranean alpine environments, and the drivers of these patterns using a comprehensive set of environmental factors across a broad temporal span. Our results show that dwarf shrubs are widely distributed in our study area and highlight a general encroachment trend over the last 60 years. Nevertheless, we found that at a fine scale, these overall trends are strongly shaped by the joint influence of the local topography, productivity, land use and micro-climate. In particular, distribution and encroachment of dwarf shrubs appear to be particularly associated with areas with harsher alpine environmental constraints and stronger resource limitation (steep and summit areas with early snowmelt and lower productivity). Thus, dwarf shrubs appear as a stress-tolerant, pioneer vegetation that is currently distributed mainly over areas that are otherwise sparsely vegetated. Moreover, our study suggests that dwarf shrubs could potentially cope with the future climatic conditions of high-altitude environments, mainly in moderate temperature increase scenarios bringing about less snow protection in winter combined with occasional heat waves and drought events in summer. On the contrary, dwarf shrubs exhibit poor competitive ability to invade grasslands compared to other woody vegetation and they are therefore likely to remain restricted to the least productive areas. Overall, our study highlights that even where shrub encroachment is widespread and appears linked to overall climatic and land-use changes, fine-scale environmental heterogeneity may strongly influence the dynamics, spatial distributions, and future responses of dwarf shrubs in the alpine vegetation mosaic.

## Acknowledgments

We are also very grateful to Francesca Napoleone, Emanuele Pelella, Livia Benedini, Giacomo Grosso and Giorgio Scarnecchia for their invaluable field and data collection assistance. We are grateful to Jaroslav Janošek for for improving the English language. Finally, the Grant to the Department of Science Roma Tre University (MIUR-Italy, Dipartimenti di Eccellenza, Articolo 1. Commi 314-337 Legge 232/2016) is gratefully acknowledged.

---

Appendix S1: land-use change in the study area

Appendix S2: grazing index validation

Appendix S3: downscaling of the climatic layer

